# DLKcat cannot predict meaningful *k*_cat_ values for mutants and unfamiliar enzymes

**DOI:** 10.1101/2023.02.06.526991

**Authors:** Alexander Kroll, Martin J. Lercher

## Abstract

The recently published DLKcat model, a deep learning approach for predicting enzyme turnover numbers (*k*_cat_), claims to enable high-throughput kcat predictions for metabolic enzymes from any organism and to capture *k*_cat_ changes for mutated enzymes. Here, we critically evaluate these claims. We show that DLKcat predictions become positively misleading for enzymes with less than 60% sequence identity to the training data, performing worse than simply assuming a mean *k*_cat_ value for all reactions. Furthermore, DLKcat’s ability to predict mutation effects is much weaker than implied, capturing only 3% of the experimentally observed variation across mutants not included in the training data. These findings highlight significant limitations in DLKcat’s generalizability and its practical utility for predicting *k*_cat_ values for novel enzyme families or mutants, which are crucial applications in fields such as metabolic modeling.

## Main text

The turnover number *k*_cat_ quantifies the catalytic efficiency of enzymes. As experimental *k*_cat_ estimates are expensive and time-consuming, it is desirable to develop computational pipelines that can predict turnover numbers of arbitrary enzymes from easily accessible features.

Advances in deep learning have now put such predictions into reach^1^. In a recent publication, Li et al.^2^ described DLKcat, a general deep learning model for *k*_cat_ predictions. The authors state that their approach facilitates “high-throughput *k*_cat_ prediction for metabolic enzymes from any organism merely from substrate structures and protein sequences.” Furthermore, they claim that “DLKcat can capture *k*_cat_ changes for mutated enzymes”. Here, we show that DLKcat predictions are accurate only for enzymes that are highly similar to proteins used for training and become positively misleading for enzymes without close homologs in the training data. We further show that DLKcat’s mutant predictions – all of which were made for enzymes highly similar to training data – are much less accurate than implied by the DLKcat publication, capturing only 3% of the experimentally observed variation across mutants not included in the training data.

The performance of machine learning predictions for enzyme features depends strongly on the sequence similarity between a target enzyme and enzymes in the training set^3^, consistent with the widely held notion that enzymes with more similar amino acid sequences are more likely to be functionally similar ^5^. Accordingly, it is likely much easier to predict unknown *k*_cat_ values for enzymes used in model training than to make predictions for enzymes that have no close homologs with known kinetic constants. More than two-thirds (67.9%) of the enzymes in the DLKcat test set are also included in the training data, and an additional 23.3% have amino acid sequences that are at least 99% identical to sequences in the training data. This extremely high similarity between training and test data constitutes a central problem in the construction of DLKcat^6^, given that its authors aimed to generate a prediction model that generalizes well to "enzymes from any organism” – i.e., to proteins with amino acid sequences that may often differ by more than a few percent from those in the training data.

The red line in **Fig. 1** shows how DLKcat’s prediction quality depends on the sequence identity between an enzyme in the test dataset and the most similar enzymes used for training. The figure shows sliding window estimates of the coefficient of determination, *R*^2^, which is a widely used measure for prediction quality (see equation (2) in Li et al. ^2^). *R*^2^ =1 indicates perfect predictions. As seen from **Fig. 1**, DLKcat’s coefficients of determination are negative for maximal sequence identities between test and training data below 60%. This indicates that when no close homologs have been used for training, DLKcat predictions are typically worse than simply assuming the same mean *k*_cat_ value for all reactions, which corresponds to *R*^2^ =0.

**Figure 1.**
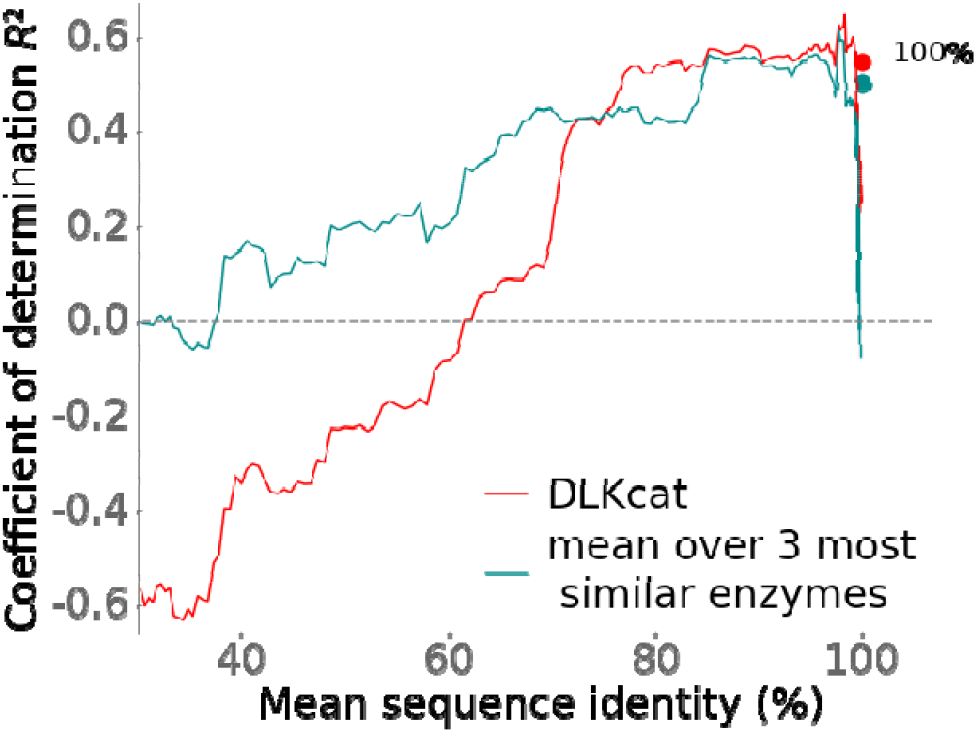
DLKcat predictions become reasonable only when closely related enzymes were used for training (max. sequence identity > 70%) and are barely better than simple *k*_cat_ averages even when the same enzyme was used for training. The curves are coefficients of determination *R*^2^, calculated in sliding windows of size *n*=100 across sequences in the test set ordered by the maximal sequence identity between individual test enzymes and all sequences in the training data. Position on the x-axis indicates the mean across the window. DLKcat predictions are shown in red. For comparison, the cyan line shows the geometric mean of *k*_cat_ values, calculated over the three most similar enzymes in the training set (without considering the catalyzed reactions or substrates; *R*^2^ =0.420 of the mean approach vs. *R*^*2*^ = 0.445 for DLKcat, calculated across the complete test dataset, *N* = 1687). The filled circles at the top right are for test datapoints with enzymes already used for training (100% max. sequence identity, *N*=1143); these were not included in the sliding windows.

When attempting to predict the effects of mutations, DLKcat is even less able to generalize beyond the proteins used for training. To support DLKcat’s ability to predict mutation effects, Li et al. calculated a combined Pearson correlation across the measured and predicted kcat values for mutants of different enzyme-substrate pairs. However, due to systematic differences across enzyme-substrate pairs, this design would lead to strong correlations even for a prediction model that simply outputs the wildtype *k*_cat_ for all mutants of a given enzyme. To assess mutation effects, we need to focus instead on the differences between the *k*_cat_s of closely related variants of the same enzyme. Thus, to facilitate a global analysis of mutant prediction quality, we grouped all mutants for the same enzyme-substrate pair and scaled the kcat values for each group to z-scores (see **Fig. 2** caption for details). 385 out of 744 mutants in the test dataset were also part of the training set, so they are not informative with respect to DLKcat’s ability to extrapolate beyond the proteins used for training. Except for five enzymes, all others have at least 99% amino acid sequence identity (*N* = 354) to sequences in the training set. Although these mutants are almost identical to proteins used for training, DLKcat cannot successfully predict the mutation effects on *k*_cat_, indicated by a negative coefficient of determination (*R*^*2*^ = −0.47; Fig. 2a).

**Figure 2.**
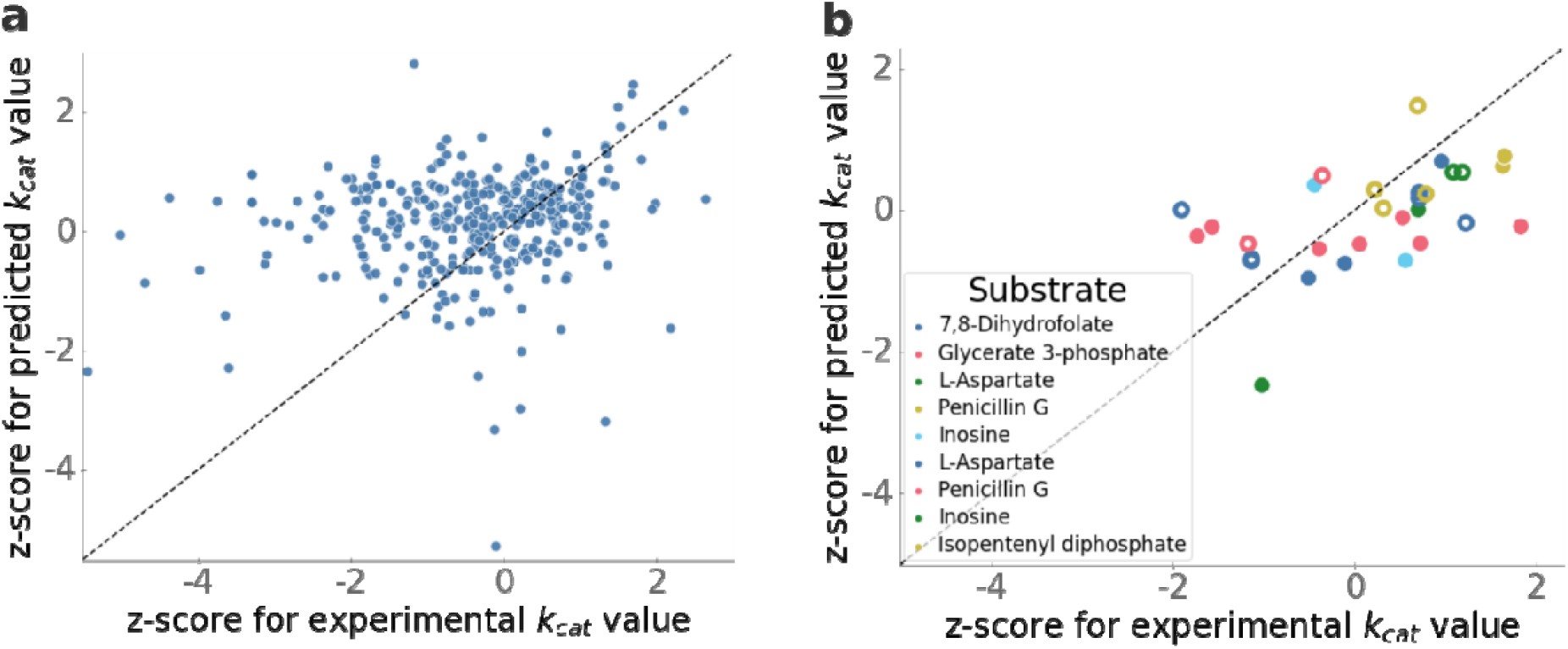
DLKcat predicts only a small fraction of *k*_cat_ variation due to mutations even for mutants that are highly similar to proteins used for training. **(a)** For all mutants in the test set with maximum sequence identities above 99% and below 100% compared to sequences in the training set, we compared predicted to experimentally observed values. To allow a global analysis of mutation effects on *k*_cat_, we scaled the log10-transformed *k*_cat_ values to z-scores as follows. For each enzyme-substrate pair, we calculated the mean and standard deviation across all measured *k*_cat_ for all other enzymes with the same substrate and with sequence identities above 98%; from each measured and each predicted *k*_cat_ for this subset, we then subtracted the mean and divided the result by the standard deviation. **(b)** For all mutants in Fig. 3c of Ref. 2, we compared predicted and experimentally observed changes in *k*_cat_ due to the mutations. Only data points from the test dataset are shown. Colors indicate different enzyme-substrate pairs. Open circles are mutants identical to enzymes in the training data, while solid dots are mutants with between 99.4% and 99.8% sequence identity to enzymes in the training data. We scaled the log10-transformed *k*_cat_ values to z-scores in a similar manner as described for panel (a). For each enzyme-substrate pair, we calculated mean and standard deviation across all measured *k*_cat_ for the wild-type and the different mutants in the complete dataset; from each measured and each predicted *k*_cat_ for this enzyme-substrate pair, we then subtracted the mean and divided the result by the standard deviation.

Arguably the most striking result presented in Ref. 2 is its Fig. 3c, which presents a “comparison between predicted and measured *k*_cat_ values for several well-studied enzyme-substrate pairs with rich experimental mutagenesis data”. The panel is augmented with a Pearson correlation coefficient *r*=0.94 and corresponding p-value < 10^−91^, suggesting that DLKcat can accurately predict the effects of mutations on *k*_cat_. However, only 15.6% of the data in this panel had been set aside for testing, while 71.9% of the datapoints shown were part of the training data. The remaining 12.5% of the datapoints were part of the validation data, which was used to select the hyperparameters of the prediction model. Any meaningful analysis of DLKcat’s abilities must be restricted to data from the test set, as done in **Fig. 2b**, which again uses z-scaling to provide a more appropriate presentation of the Fig. 3c data. 14 of the 30 mutant enzymes in the test set had also been used for training, and each of the remaining 16 mutant amino acid sequences is at least 99.4% identical to a sequence in the training set. Despite this close relationship between test and training enzymes, DLKcat predicts only a small fraction of the variation in *k*_cat_ due to mutations: when calculating a coefficient of determination across the z-scores to focus on mutation effects (rather than baseline effects of the different enzyme-substrate pairs), one obtains a low coefficient of determination, *R*^2^=0.162. When restricting this analysis to the 16 mutants that were not in the training data, the coefficient of determination is further reduced to *R*^*2*^ =0.030. Thus, for previously unseen mutants of the enzyme-substrate pairs in Li *et al*.’s Fig. 3c, DLKcat predicts only 3% of the variance relative to the wildtype *k*_cat_, even though these mutants are highly similar to wild-type and/or mutant enzymes in the training data.

In sum, while Li et al. claim that DLKcat can predict *k*_cat_ for metabolic enzymes from any organism and can capture *k*_cat_ changes for mutated enzymes, the above analyses demonstrate that these claims are inconsistent with a careful analysis of the data in Ref. 2. For enzymes without close homologs in the training data (<60% sequence identity), DLKcat predictions are positively misleading (*R*^*2*^ <0, **Fig. 1**), and DLKcat makes no meaningful predictions for mutants not included in the training set. Thus, DLKcat provides little benefit when predicting turnover numbers for enzyme families and mutants not already characterized kinetically, which arguably constitute the most important use cases in most applications, including the parameterization of enzyme-constrained genome-scale metabolic models (ecGEMs). This disappointing performance is likely rooted in the construction of the DLKcat datasets, which did not challenge the model to learn to predict *k*_cat_ values for enzymes dissimilar to those used for training.

## Methods

To reproduce the DLKcat model and to make predictions for the corresponding test set, we downloaded the code provided on GitHub by Li et al.^2^. To calculate the maximal amino acid sequence identity between each enzyme in the test set and all enzymes in the training set, we used the Needleman-Wunsch algorithm implemented in the EMBOSS software package ^7^.

## Code and data availability

All software was coded in Python 3. The code used to generate the results of this paper, in the form of Jupyter notebooks, as well as all datasets, are available from https://github.com/AlexanderKroll/DLkcat_Matters_Arising.

## Acknowledgements

We are grateful to the authors of Ref. 2 for comments on an earlier draft. Computational support and infrastructure were provided by the “Centre for Information and Media Technology” (ZIM) at Heinrich Heine University Düsseldorf (Germany). This work was funded by the Deutsche Forschungsgemeinschaft (DFG, German Research Foundation) through CRC 1310, and, under Germany’s Excellence Strategy, through grant EXC 2048/1 (Project ID: 390686111).

## Author contributions

AK & MJL conceived of the study, performed the analyses, interpreted the results, and wrote the manuscript.

